# Interaction Between Host MicroRNAs and the Gut Microbiota in Colorectal Cancer

**DOI:** 10.1101/192401

**Authors:** Ce Yuan, Michael Burns, Subbaya Subramanian, Ran Blekhman

## Abstract

**Background:** Although variation in gut microbiome composition has been linked with colorectal cancer (CRC), the factors that mediate the interactions between CRC tumors and the microbiome are poorly understood. MicroRNAs (miRNAs) are known to regulate CRC progression and patient survival outcomes. In addition, recent studies suggested that host miRNAs can also regulate bacterial growth and influence the composition of the gut microbiome. Here, we investigated the association between miRNAs expression in human CRC tumor and normal tissues and the microbiome composition associated with these same tissues.

**Method:** We sequenced the small RNAs from patient-matched tumor and normal tissue samples collected from 44 human CRC patients performed an integrated analysis with microbiome taxonomic composition data from these same samples. We then interrogated the functions of the bacteria correlated with miRNAs that were differentially expressed (DE) between tumor and matched normal tissues, as well as the functions of miRNAs correlated with bacterial taxa that have been previously associated with CRC, including *Fusobacterium, Providencia, Bacteroides, Akkermansia, Roseburia, Porphyromonas, and Peptostreptococcus.*

**Results:** We identified 76 miRNAs as DE between CRC and normal tissue, including known oncogenic miRNAs miR-182, miR-503, and miR-17∼92. These DE miRNAs were correlated with the relative abundance of several bacterial taxa, including Firmicutes, Bacteroidetes, and Proteobacteria. Bacteria correlated with DE miRNAs were enriched with distinct predicted metabolic categories. Additionally, we found that miRNAs correlated with CRC-associated bacteria are predicted to regulate targets that are relevant for host-microbiome interactions, and highlight a possible role for miRNA-driven glycan production in the recruitment of pathogenic microbial taxa.

**Conclusions:** Our work characterized a global relationship between microbial community composition and miRNA expression in human CRC tissues. Our results support a role for miRNAs in mediating a bi-directional host-microbiome interaction in CRC. In addition, we highlight sets of potentially interacting microbes and host miRNAs, suggesting several pathways that can be targeted via future therapies.

## Introduction

The colon microenvironment hosts trillions of microbes, known as the gut microbiome. A healthy microbiome helps maintain colon microenvironment homeostasis, immune system development, gut epithelial function, and other organ functions [1–5]. Although many factors impact the composition of the gut microbiome, the overall functional profiles remain stable over time [6, 7]. Nevertheless, changes in the taxonomic and functional composition of the microbiome have been implicated in many diseases, including colorectal cancer (CRC) [8–11]. Although the association between microbiome alterations and disease processes has been extensively demonstrated, the directionality, as well as the mediators of the host-microbiome interaction, remain unclear.

Diet has been independently associated with both the gut microbiome and CRC. For example, the western diet (characterized by low fiber and high protein, fat, and sugar) affects gut microbiome composition in humanized mice, whereby mice fed with western diet have increased Firmicutes and decreased Bacteroidetes relative abundance [12, 13]. The same western diet has also long been considered as a risk factor for developing CRC [14–16]. Using an animal model of CRC, Schulz *et al.* demonstrated that the high-fat diet (HFD) exacerbates CRC progression; however, treating animals with antibiotics blocks HFD-induced CRC progression [17]. This suggests that diet can drive microbiome composition change in the gut as a precursor to CRC development.

Recent studies have found that host genetic variation can affect microbiome composition. For example, a polymorphism near the *LCT* gene, which encodes the lactase enzyme, is associated with the abundance of *Bifidobacterium* in the gut microbiome, and *Christensenellaceae* family are shown to be heritable, with a higher similarity between monozygotic than dizygotic twins. [18–23]. This could explain uni-directional host-to-microbiome interaction, especially in CRC, where genetic mutations in host cells are common [24, 25]. Interestingly, in a genetic mutation model of intestinal tumors, germ-free animals developed significantly fewer tumors in the small intestine [26]. Although the finding is limited to the small intestine, the trend shows that CRC development partially depends on the microbiome. In an animal model of colitis-associated CRC, Uronis *et al.* showed that germ-free mice exhibit normal histology and do not develop tumors, compared to 62% for conventionalized mice that have developed tumors (n=13) [27]. Additionally, Ridaura *et al.* demonstrated that when transplanting the microbiome from twins discordant for obesity to germ-free mice, the mice that received the obesity-associated microbiome developed an obese phenotype while the mice that received lean microbiome did not [13]. Clearly, this evidence suggests that host genetics only partially explains host-microbiome interaction.

A recent report demonstrated that a fecal microRNAs (miRNAs) can shape the composition of the gut microbiome [28], indicating a mechanism by which host cells can regulate the microbial community. In CRC, several miRNAs, such as miR-182, miR-503, and miR-17∼92 cluster, can regulate multiple genes and pathways and have been found to promote malignant transformation and disease progression [29–31]. Interestingly, studies have also found that microbiome-derived metabolites can change host gene expression, including expression of miRNAs, in the colon [32, 33]. Taken together, these results suggest a bi-directional interaction between host cells and microbes, potentially mediated through miRNA activity. However, we still know very little about the role of miRNAs in host-microbiome interaction, especially in the context of CRC. With thousands of unique miRNAs and microbial taxa present in the CRC microenvironment, it is challenging to experimentally study all possible pairwise interactions. Nevertheless, genomic characterization of both miRNA expression and microbial composition in CRC can identify potential interactions between miRNAs and microbes, which can then be used as candidates for functional inspection.

Here, we establish the relationships between miRNAs expression and microbiome composition in CRC patients. We sequenced small RNAs and integrated 16S rRNA gene sequencing data from both tumor and normal colon tissues from 44 patients (88 samples total). We explored the correlation between miRNAs and the microbiome through imputing the miRNA functional pathways and microbiome metabolic pathways *in silico* **(Supplementary Figure 1)**. To our knowledge, this is the first analysis to establish the global relationship between miRNAs expression and microbiome in CRC patients.

**Figure 1:**
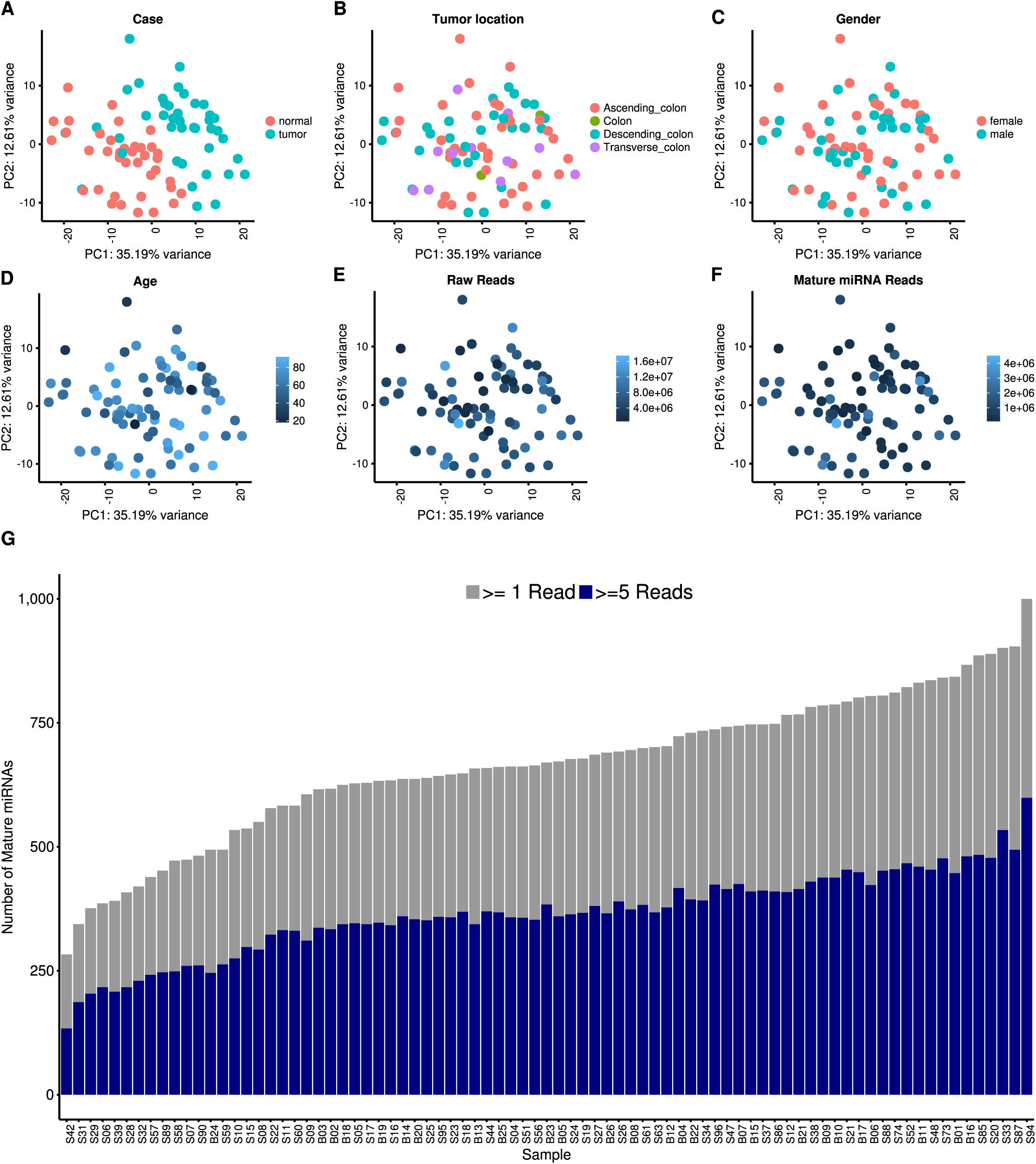
Small RNA sequencing data quality. Principle component analysis showing PC1 on the x-axis and PC2 on the y-axis. Each dot is colored by **(a)** normal/tumor, **(b)** tumor location, **(c)** patient gender, **(d)** patient age, **(e)** raw read count, and **(f)** mature miRNA mapped read count. **(g)** Bar plot of the numbers of mature miRNAs identified in each sample, with coverage over 1 read (gray) and over 5 reads (blue).

## Methods

### Tissue samples

A total of 88 matched tumor and adjacent normal tissues were collected from 44 patients by the University of Minnesota Biological Materials Procurement Network. A detailed description of sample collection was previously published [8]. Briefly, all patients provided written, informed consent. All research conformed to the Helsinki Declaration and was approved by the University of Minnesota Institutional Review Board, Protocol 1310E44403. Tissue pairs were resected concurrently, rinsed with sterile water, flash frozen in liquid nitrogen, and characterized by staff pathologists. Detailed deidentified sample metadata, including age, gender, tumor location, tumor stage, and microsatellite stability (MSS) status is available in **Supplementary Table 1**.

### 16S rRNA sequencing and sequence analysis

The 16S rRNA gene sequencing data were previously published [8]. Raw sequences were deposited at NCBI Sequence Read Archive under project accession PRJNA284355, and processed data files are available in Burns *et al.* [8]. Briefly, total DNA was extracted from approximately 100 mg of tissue. Tissues were first physically disrupted by placing the tissue in 1 mL of Qiazol lysis solution in a 65 °C ultrasonic water bath for 1–2 h. The efficiency of this approach was verified by observing high abundances of Gram-negative bacteria across all samples, including those from the phylum Firmicutes. DNA was then purified using AllPrep nucleic acid extraction kit (Qiagen, Valencia, CA). The V5-V6 region of the 16S rRNA gene was PCR amplified with multiplexing barcodes [34]. The barcoded amplicons were pooled and ligated to Illumina adaptors. Sequencing was performed on a single lane on an Illumina MiSeq instrument (paired-end). The forward and reverse read pairs were merged using the USEARCH v7 program ‘fastq_mergepairs’, allowing stagger, with no mismatches, allowed [35]. OTUs were picked using the closedreference picking script in QIIME v1.7.0 using the Greengenes database (August 2013 release) [36–38]. The similarity threshold was set at 97%, reverse read matching was enabled, and reference based chimera calling was disabled. The unfiltered OTU table used for the analysis is available in **Supplementary Table 1**.

### MicroRNA sequencing

To prepare samples for small RNA sequencing, total RNA was extracted using AllPrep nucleic acid extraction kit (Qiagen, Valencia, CA). RNA was quantified using RiboGreen fluorometric assay (Thermo Fisher, Waltham, WA). RNA integrity was then measured using BioAnalyzer 2100 (Agilent, Santa Clara, CA). Library creation and sequencing were performed by the Mayo Clinic Genome Analysis Core. Briefly, small RNA libraries were prepared using 1 ug of total RNA per the manufacturer’s instructions for NEBNext Multiplex Small RNA Kit (New England Biolabs; Ipswich, MA). After purification of the amplified cDNA constructs, the concentration and size distribution of the PCR products was determined using an Agilent Bioanalyzer DNA 1000 chip (Santa Clara, CA) and Qubit fluorometry (Invitrogen, Carlsbad, CA). Four of the cDNA constructs are pooled and the 120-160bp miRNA library fraction is selected using Pippin Prep (Sage Science, Beverly, MA). The concentration and size distribution of the completed libraries was determined using an Agilent Bioanalyzer DNA 1000 chip (Santa Clara, CA) and Qubit fluorometry (Invitrogen, Carlsbad, CA). Sequencing was performed across 4 lanes on an Illumina HiSeq 2000 instrument (paired end).

### MicroRNA sequence data processing and QC

See **Supplementary Figure 1** for an overview of the data analysis steps. Briefly, quality control of miRNA sequencing data was performed using FastQC before and after adaptor trimming with trimmomatic [39]. Then, the paired-end reads were assembled using PANDASeq and aligned to hg38 genome assembly using bowtie2 [40, 41]. Finally, the total mature miRNA counts were generated with HTseq [42]. We removed 7 samples due to low number of total raw reads (fewer than 500,000 raw reads) from the analysis. The remaining 81 samples have between 519,373 and 17,048,093 (median 6,010,361) reads per library, with an average quality score greater than 37 in all libraries. Between 66.79% and 96.14% (median 83.53%) of reads passed adapter trimming **(Supplementary Figure 2)**. Of all the reads passing adapter trimming, between 287,356 and 11,102,869 (median 3,701,487) reads are identified as concordant pairs by PANDASeq. After mapping to hg38 genome, between 18,947 and 4,499,805 (median 859,546) reads were assigned to a total of 2,588 mature miRNAs **(Supplementary Figure 3)**. Principle component analysis (PCA) showed a clear separation between tumor and normal samples (**Figure 1A**), while tumor location, gender, age, total raw reads, and total mature miRNA reads do not appear to have an impact on the data **(Figure 1B-F)**. Similarly, a PCA plots including an additional principal component did not detect clustering based on these factors (**Supplementary Figure 4**). Between 283 and 1,000 (median 670) miRNAs had coverage over 1 read, and between 134 and 599 (median 367) miRNAs had coverage over 5 reads **(Figure 1G)**. Overall the quality of our sequencing results is on par with previous studies and our previous observations [43]

**Figure 2:**
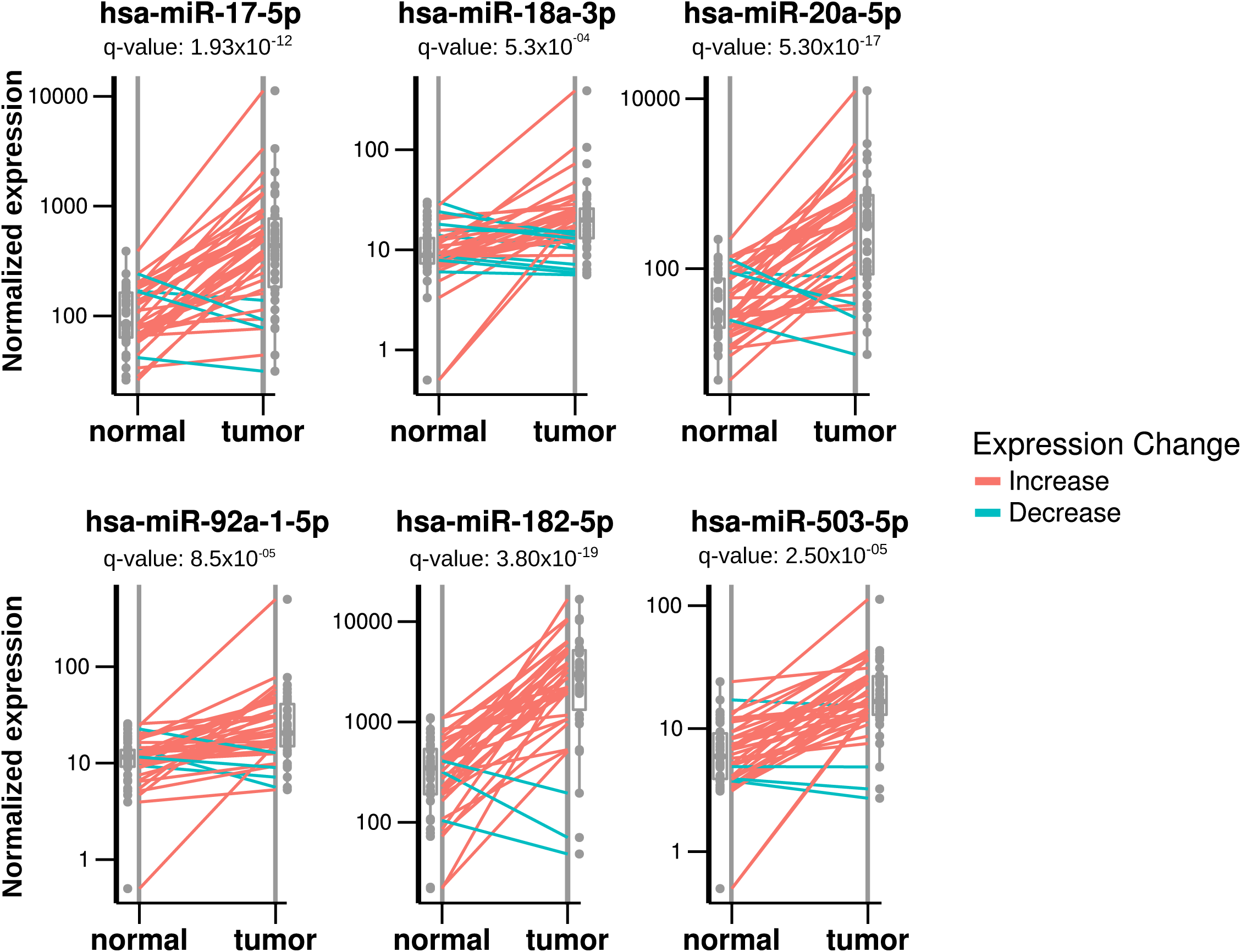
Differentially expressed miRNAs between matched normal and tumor. Boxplot and dot plot showing differentially expressed miRNAs. Each panel represents a single miRNA with normalized expression level on the y-axis. Lines connects a normal and tumor sample from the same individual, with red lines indicate higher expression level in tumor tissues and green lines indicate higher expression level in normal tissues. miR-17,-18a,-20a,92a,-182 and -503 were found to have significantly higher expression levels in tumor tissues.

**Figure 3:**
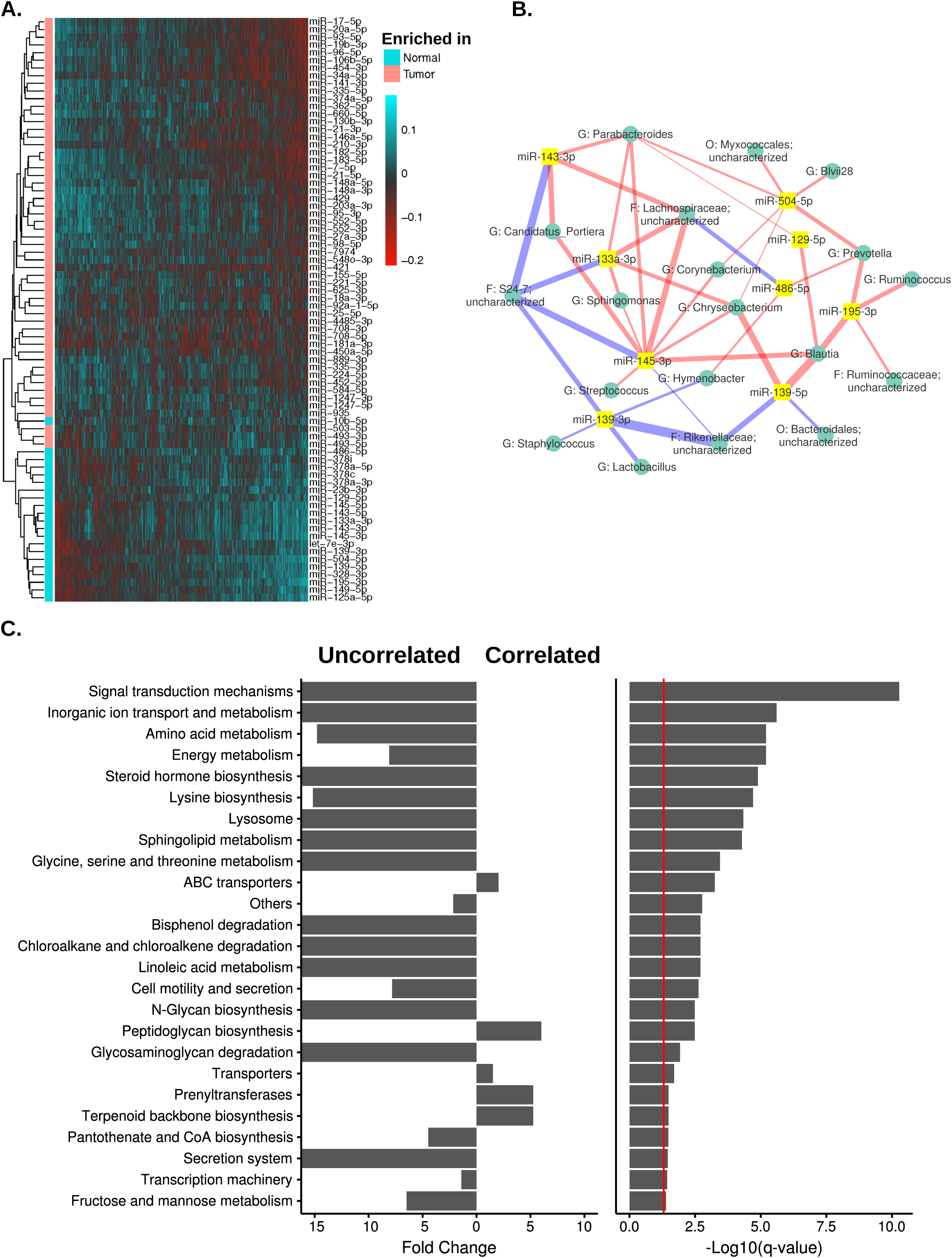
Bacteria significantly correlated with DE miRNAs. **(a)** Heatmap showing bacterial genera (in columns) significantly correlated with the DE miRNAs (in rows). Red indicates negative correlations and green indicates positive correlations. **(b)** Interaction network showing the ten most significantly DE miRNAs and their correlated bacteria (showing bacteria with relative abundance > 0.1% and correlation pseudo p-value ≤ 0.05). Edge thickness represents the magnitude of the correlation, with blue indicating negative correlation while red indicating positive correlation. **(c)** Metabolic pathway (KEGG) enrichment of microbiome correlated and uncorrelated with DE miRNAs. The bar graph (left panel) shows the fold enrichment for each group. FDR corrected p-value from Wilcoxon Rank Sum test (on a negative log10 scale) are shown on the right panel. The solid red line indicates q-value of 0.05.

**Figure 4:**
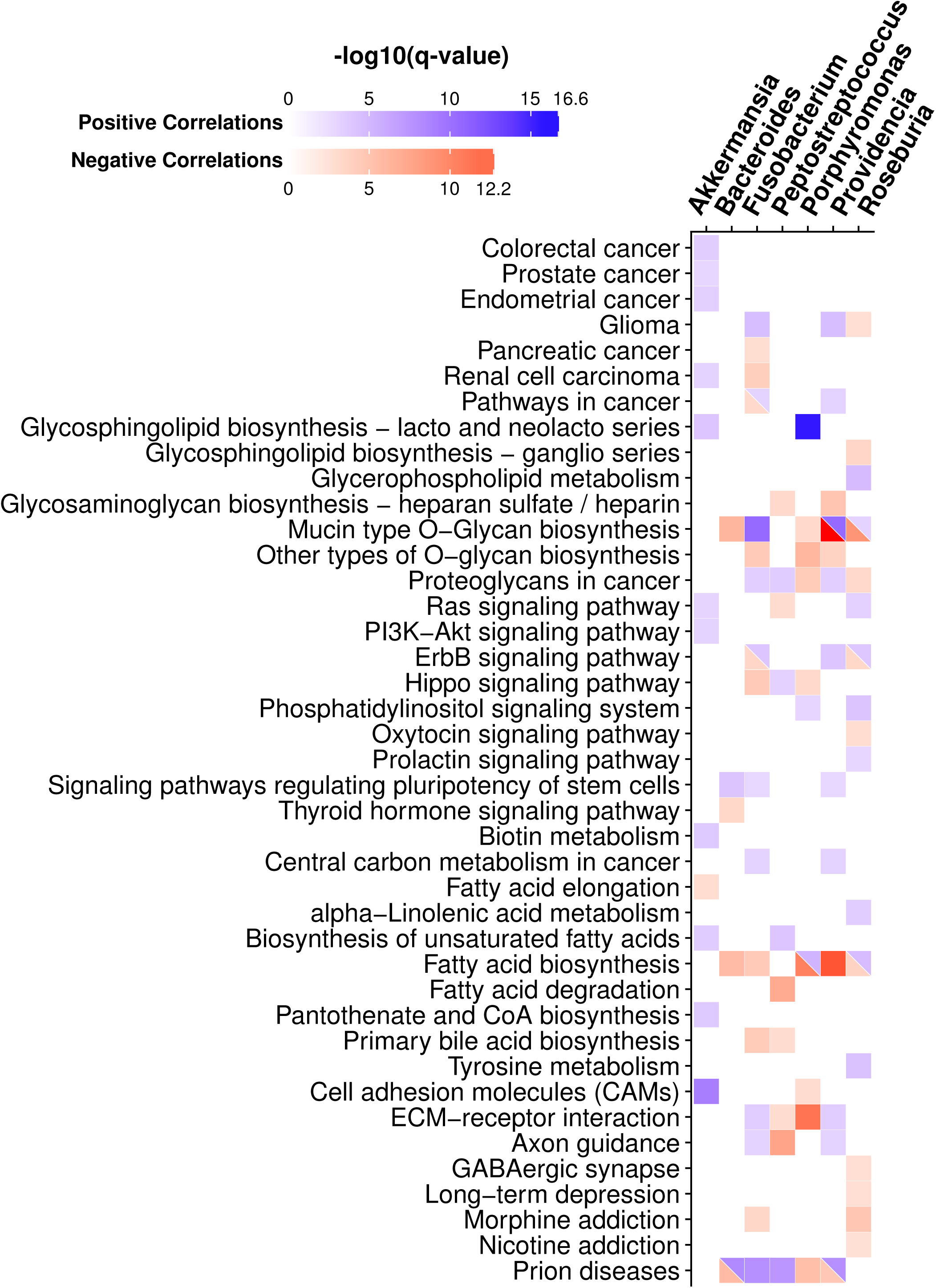
miRNA target pathways correlated with CRC-associated bacteria. The heat map shows the predicted pathways of miRNAs (rows) correlated with CRC-associated bacteria (columns) with q-value < 0.01 (modified Fisher’s Exact Test; FDR corrected). Positive correlations are shown in blue and negative correlations are shown in red. The color intensity is shown in -log10 scale of FDR corrected p-value from modified Fisher’s exact test generated by miRPath, with darker color indicating lower q-value.

### MicroRNA differential expression and correlation analysis

We identified differentially expressed (DE) miRNAs between tumor and normal samples using the DESeq2 package (1.10.1) in R (version 3.2.3) [44]. Raw miRNA counts were filtered to include miRNAs with ≥ 1 read in ≥ 80% of the samples. The remaining 392 miRNAs were then used for DESeq2 analysis. We define DE miRNAs as showing fold change over 1.5 with false discovery rate (FDR) adjusted p-value (q-value) < 0.05. We performed correlation analysis using Sparse Correlations for Compositional data (SparCC) at the genus level for bacteria [45]. To increase the accuracy of estimation, we performed 20 iterations for each SparCC procedure. We calculated pseudo p-values using 100 random permutations. Significant correlations were defined as r over 0.05 (or less than -0.05) with a pseudo p-value ≤ 0.05 [8]. Heat maps of the correlation were generated in R using the pheatmap package. We performed hierarchical clustering for both column and rows with average linkage method using Pearson’s correlation. We included bacteria with significant correlations with DE miRNAs to perform metagenomic prediction using PICRUSt v1.0.0 [46]. We included miRNAs with significant correlations with CRC associated genera (*Fusobacterium, Providencia, Bacteroides, Akkermansia, Roseburia, Porphyromonas, and Peptostreptococcus)* to perform pathway enrichment analysis using miRPath v.3 [47, 48]. We generated network visualization of miRNA-microbe using Cytoscape v3.5.1

## Results

### MicroRNAs differentially expressed in tumor tissues

Before performing differential expression (DE) analysis, we performed extensive quality control of the miRNA data. Our results indicate that miRNA expression is not strongly affected by tumor location, patient gender, patient age, and read coverage, and shows a clear clustering of miRNA data by tumor and normal samples (see **Figure 1** and Methods above). To identify small RNAs that are DE between tumor and normal samples, we performed DE analysis using DESeq2 (see Methods). A total of 76 DE miRNAs were identified, with 55 up-regulated and 21 down-regulated in tumor tissues compared to normal tissues (p-value < 0.05 after FDR correction). A full list of DE miRNAs is available in **Supplementary Table 3**. DE miRNAs with higher expression levels in tumor tissues include miR-182, miR-183, miR-503, and miR-17∼92 cluster miRNAs **(Figure 2; Supplementary Table 3)**, all consistent with our previous reports [29, 49]. These miRNAs have all been previously shown to contribute to CRC disease progression; for example, miR-182 and miR-503 were found to cooperatively target *FBXW7* and contribute to CRC malignant transformation and progression, and were also predictive of patient survival [29]. The miR-17∼92 cluster regulates multiple tumor-suppressive genes in CRC and other cancers [50]. In addition, miR-1, miR-133a, and miR-448 **(Supplementary Table 3)** were observed at higher levels in normal tissues compared with matched tumor tissues, also in agreement with previous reports [49, 51].

### Predicted functions of microbiome taxa correlated with DE miRNAs

To investigate the relationship between individual miRNA and microbiome in CRC tumor samples, we performed correlation analysis using SparCC. SparCC is developed specifically to analyze compositional genomic survey data, such as 16S rRNA gene sequencing and other types of high-throughput sequencing data [45]. Hierarchical clustering revealed several clusters of significantly correlated miRNAs and bacterial taxa **(Supplementary Figure 5)**. To further investigate the relationship between miRNAs and microbiome in CRC we selected bacteria significantly correlated with the DE miRNAs **(Figure 3A)**. The correlations clearly show a distinct pattern based on the enrichment of miRNAs, even though the correlation analysis is performed only in tumor samples. We then built a network visualizing the relationship between the top 10 DE miRNAs and their significantly correlated bacteria **(Figure 3B)**. The correlation network shows a highly-interconnected relationship between these miRNAs and bacteria. Interestingly, *Blautia*, a genus previously found to have lower abundance in tumor samples, is negatively correlated with miR-20a, miR-21, miR-96, miR-182, miR-183, and miR-7974, which are all miRNAs with high expression levels in tumor tissues. *Blautia* is also positively correlated with the expression level of miR-139, which is a miRNA with high expression levels in normal tissues. Experimental validations are required to investigate the correlations.

We then analyzed the predicted functional composition of the microbiome data and investigated correlations with miRNAs **(Figure 3C).** We hypothesized that if miRNAs selectively affect the growth of certain bacteria, then bacteria correlated with DE miRNAs are likely to represent functional differences between tumor and normal tissues, while the uncorrelated bacteria would not. Using the PICRUSt v.1.0.0 software, we generated the predicted functional profiles of the correlated and uncorrelated bacteria by assigning pathways and enzymes using the Kyoto Encyclopedia of Genes and Genomes (KEGG) database. A total of 25 pathways have significantly altered enrichment (two-sided Wilcoxon signed rank test with FDR corrected p-value < 0.05; **Figure 3C**). Interestingly, several metabolic pathways and signaling pathways, including signal transduction, amino acid metabolism, energy metabolism, and linoleic acid metabolism, were all enriched in the uncorrelated group, suggesting increased metabolic processes in this group. For bacteria significantly correlated with DE miRNAs, however, pathways related to transporters, peptidoglycan, and terpenoid backbone biosynthesis have significant enrichment.

### Predicted functions of miRNAs correlated with CRC associated bacteria

To investigate the function of miRNAs correlated with CRC-associated bacteria, we focused on bacteria genera previously associated with CRC, including *Fusobacterium, Providencia, Bacteroides, Akkermansia, Roseburia, Porphyromonas, and Peptostreptococcus* [8, 52–55]. We hypothesized that if these bacteria affect CRC through modulating miRNA expression, then miRNAs that are significantly correlated with the bacteria are expected to show enrichment in cancer-related genes and pathways. A list of miRNAs significantly correlated with these bacteria is available in **Supplementary Table 4**. We separated these miRNAs into groups with positive correlation and negative correlation with each bacteria independently. Then, using the miRPath v.3 software, we predicted the functions of miRNAs by assigning pathways to the miRNA targets using the KEGG database **(Supplementary Table 5)**. We visualized the pathways with q-value < 0.01 (modified Fisher’s Exact Test; FDR corrected) in **Figure 4**.

Our results show that *Akkermansia* is the only taxon correlates with miRNAs associated with “colorectal cancer” pathway. *Fusobacterium, Providencia* and *Roseburia* correlate with miRNAs associated with cancer-related pathways, including “Glioma”, “Pancreatic cancer”, “Renal cell carcinoma” and “Pathways in cancer”. Interestingly, glycan-related pathways, including “mucin-type O-glycan biosynthesis”, “Other O-Glycan biosynthesis”, “Glycosaminoglycan biosynthesis - heparan sulfate/heparin”, and “Proteoglycans in cancer”, have correlations with all bacteria genera analyzed, except for *Akkermansia*. This finding corresponds to a previous study showing that *Fusobacterium nucleatum* infection stimulates mucin secretion *in vitro* [56]. Additionally, *Fusobacterium nucleatum* binds to specific Gal-GalNAc, which is expressed by CRC tumors, through the *Fap2* protein [57]. *Porphyromonas gingivalis* were shown to induce a proteoglycan, syndecan-1, shedding in oral epithelial cells [58]. However, the role of the bacteria and glycan interaction is not clear in the context CRC. Cell signaling pathways previously implicated in CRC, such as Ras, PI3K/Akt, ErbB and Hippo pathways are also correlated with these bacteria [59–62].

## Discussion

Although there is a known association between gut microbiome composition change and CRC [8–11], the potential mediators of this relationship remain unclear. One potential mediator is host genetics, and specifically CRC tumor mutational profiles [26, 27]. Additional evidence indicates that miRNAs can mediate host-microbiome interactions in CRC [28]. Here, we presented the first integrated analysis of miRNA expression and gut microbiome profiles in CRC patients. Our data show a highly interconnected correlation network between miRNA expression and the composition of the microbiome and supports the role for miRNAs in mediating host-microbiome interactions.

Active interactions between host and the microbiome in CRC has been previously observed, leading to the proposition that pathogenic “passenger” bacteria colonizing tumor tissue could lead to exacerbated tumor progression [63]. In our analysis, we focused on potential “passenger” bacteria, including *Fusobacterium, Providencia, Bacteroides, Akkermansia, Roseburia, Porphyromonas, and Peptostreptococcus*. *Fusobacterium* includes several pathogenic species, and are implicated in dental disease, infections, and CRC [64–66]. Similarly, *Providencia* has also been implicated in gastrointestinal infections [8, 67–69]. The mechanism of *Fusobacterium* in promoting CRC tumorigenesis and progression has been investigated. It activates the *Wnt/β-catenin* signaling pathway through *FadA* protein which binds to the E-cadherin protein on the intestinal epithelial cells (IECs), thus promoting cell proliferation [64]. Several mechanisms could explain this observation. One possibility is that bacteria can infiltrate the intestinal epithelial barrier after certain pathogenic bacteria cleaving the E-cadherin [64, 70]. This could lead to an increased inflammatory response in the colon microenvironment, and the inflammation can lead to DNA damage and contribute to disease progression [63, 64]. Another potential mechanism is that bacteria can directly cause mutations in IECs through virulence proteins. Several of these virulence proteins were found in *Escherichia coli* and *Helicobacter pylori* [71, 72], and results indicate that these virulence factors may be enriched in the CRC microbiome, especially in *Fusobacterium* and *Providencia* [8]. However, it is unclear if these bacteria produce virulence proteins that can directly cause DNA damage, and further investigation is required to elucidate this mechanism.

The *Wnt/β-catenin* pathway activation by *Fusobacterium* can lead to upregulation of numerous genes related to CRC [73–75]. One such gene, *MYC*, is a transcription factor that targets multiple genes related to cell proliferation, cell cycle, and apoptosis. The miR-17∼92 cluster is a known target of *MYC* and has oncogenic properties in several cancer types [31, 50, 76, 77]. Interestingly, butyrate, a short-chain fatty acid (SCFA) produced by members of the microbiome, diminishes *MYC-*induced miR-17∼92 overexpression in CRC *in vitro* through its function as histone deacetylase inhibitor [32]. Studies in CRC have consistently found low fecal butyrate levels as well as a reduced relative abundance of butyrate-producing bacteria, such as Firmicutes phylum [32, 52, 78]. One potential explanation is in CRC, the DE miRNAs can affect the growth of certain microbes, which eventually outcompete other species and form a biofilm on tumor tissues [28]. Indeed, our data shows several enriched bacterial nutrient biosynthesis and metabolism pathways in the microbes uncorrelated with DE miRNAs, but not in the correlated group. Interestingly, pathways in bacterial cell motility and secretion are also enriched among uncorrelated bacteria, suggesting that, in addition to promoting bacterial growth, certain miRNAs may be involved in recruiting bacteria to tumor tissues. This may also provide a possible explanation for the observed difference in alpha diversity of tumor microbiomes [8, 79, 80].

In our analysis of the functions of miRNAs correlated with selected bacteria known to have associations with CRC, the prion diseases, glioma, and morphine addiction pathways found to be enriched in our analysis do not immediately seem related to cancer (**Figure 4**). Upon further investigation of miRNAs gene targets in these pathways, we found that several genes included in the pathways may have relevant functions in cancer. For example, Mitogen-activated Protein Kinase (*MAPK*) is central to cell proliferation and survival; Interleukin-6 (*IL6*) and Interleukin-beta (*IL1β*) are cytokines involved in inflammation; Protein Kinase A (*PKA*) is important in regulating nutrient metabolism; Bcl-2-associated X protein (*BAX*) is a tumor suppressor gene; and Prion protein (*PRNP*) are known to have a significant role in regulating immune cell function [81–83].

A recent study has suggested an additional mechanism affecting host-microbiome interactions that may promote CRC tumorigenesis and progression [57, 84]. Abed *et al.* showed that *Fap2* produced by *Fusobacterium* binds to glycan produced by CRC to attach to the tumor tissue [57]. Interestingly, glycan biosynthesis pathways were enriched in targets of the miRNAs correlated with CRC-associated bacteria. The increased glycan production may increase recruitment of certain bacteria, such as *Fusobacterium*, to the tumor location. This result highlights a novel potential mechanism for miRNAs, through regulating glycan biosynthesis, to attract specific microbes to the tumor microenvironment, and thus impact tumor development. Interestingly, mucin-type O-glycan biosynthesis pathway is enriched in miRNAs positively correlated with *Fusobacterium* but negatively correlated with *Bacteroides* and *Porphyromonas*. This suggests that these bacteria may have different mechanisms of attachment to the mucosal surface due to different ability to bind to O-glycan [85]. Additional studies are required to test the association between *Fusobacterium*, tumorigenesis, and miRNA-driven glycan production.

It is important to note that our study uses 16S rRNA gene sequencing to characterize microbiome taxonomic composition and computationally predicted pathway composition using PICRUSt v1.0.0 [46]. Although this method is widely used, metagenomics shotgun sequencing can be more accurate and informative in understanding the functional makeup of a microbial community. Similarly, to impute miRNA functional profiles, we used an *in silico* prediction method, miRPath [46, 47]. While these two methods have both been rigorously tested and validated with experimental data, the results remain predictions and may not represent the real biological system [46, 47]. Another limitation of our approach is that it identifies correlations and not causal relationships. Nevertheless, this approach allows us to generate a microbiome- and miRNA transcriptome-wide characterization of potential interactions, which shed light on potential new mechanisms of host-microbiome interactions. In addition, we highlight candidates for potentially interacting host miRNAs and microbial taxa, which can be validated in model organism studies that will address causality.

## Conclusions

Our analysis, together with evidence from previous studies, suggests that miRNAs likely mediate host-microbiome interaction in CRC. We identify potential novel mechanisms that mediate this interaction and may have a role in CRC tumorigenesis, including a possible role for miRNA-driven glycan production in the recruitment of pathogenic microbial taxa. The interactions identified here could be a direct target for developing therapeutic strategies that can benefit CRC patients. Follow-up studies in model organisms are warranted to assess the causal role of individual microbes and miRNAs in CRC.

## Acknowledgements

The authors thank the members of the Blekhman and the Subramanian lab for helpful discussions. We thank Dr. Anne Sarver of Subramanian lab for developing miRNA analysis pipeline. We also thank the Medical Genome Facility Genome Analysis Core at Mayo Clinic (Rochester, MN) for performing library prep and sequencing. This work was carried out, in part, using computing resources at the Minnesota Supercomputing Institute.

## Author contributions

CY, SS, and RB developed the concept, CY carried out data analysis with assistance from MB. CY, SS, and RB wrote the manuscript.

## Funding

C.Y. is supported by Norman Wells Memorial Colorectal Cancer fellowship and Healthy Foods Healthy Lives Institute Graduate and Professional Student Research Grant. This work is supported the Randy Shaver Cancer Research and Community Fund (R.B.), Institutional Research Grant #124166-IRG-58-001-55-IRG53 from the American Cancer Society (R.B.), and a Research Fellowship from The Alfred P. Sloan Foundation (R.B.).

## References

1. Abdollahi-Roodsaz S, Abramson SB, Scher JU: The metabolic role of the gut microbiota in health and rheumatic disease: mechanisms and interventions. Nat Rev Rheumatol 2016, 12:446–455.

2. Belkaid Y, Hand TW: Role of the microbiota in immunity and inflammation. Cell 2014, 157:121–141.

3. Candon S, Perez-Arroyo A, Marquet C, Valette F, Foray A-P, Pelletier B, Milani C, Ventura M, Bach J-F, Chatenoud L: Antibiotics in early life alter the gut microbiome and increase disease incidence in a spontaneous mouse model of autoimmune insulin-dependent diabetes. PLoS ONE 2015, 10:e0125448.

4. Vangay P, Ward T, Gerber JS, Knights D: Antibiotics, pediatric dysbiosis, and disease. Cell Host Microbe 2015, 17:553–564.

5. Tremaroli V, BÄckhed F: Functional interactions between the gut microbiota and host metabolism. Nature 2012, 489:242–249.

6. David LA, Maurice CF, Carmody RN, Gootenberg DB, Button JE, Wolfe BE, Ling AV, Devlin AS, Varma Y, Fischbach MA, Biddinger SB, Dutton RJ, Turnbaugh PJ: Diet rapidly and reproducibly alters the human gut microbiome. Nature 2014, 505:559–563.

7. Faith JJ, Guruge JL, Charbonneau M, Subramanian S, Seedorf H, Goodman AL, Clemente JC, Knight R, Heath AC, Leibel RL, Rosenbaum M, Gordon JI: The long-term stability of the human gut microbiota. Science 2013, 341:1237439.

8. Burns MB, Lynch J, Starr TK, Knights D, Blekhman R: Virulence genes are a signature of the microbiome in the colorectal tumor microenvironment. Genome Med 2015, 7:55.

9. Nakatsu G, Li X, Zhou H, Sheng J, Wong SH, Wu WKK, Ng SC, Tsoi H, Dong Y, Zhang N, He Y, Kang Q, Cao L, Wang K, Zhang J, Liang Q, Yu J, Sung JJY: Gut mucosal microbiome across stages of colorectal carcinogenesis. Nat Commun 2015, 6:8727.

10. Wang T, Cai G, Qiu Y, Fei N, Zhang M, Pang X, Jia W, Cai S, Zhao L: Structural segregation of gut microbiota between colorectal cancer patients and healthy volunteers. ISME J 2012, 6:320–329.

11. Shen XJ, Rawls JF, Randall T, Burcal L, Mpande CN, Jenkins N, Jovov B, Abdo Z, Sandler RS, Keku TO: Molecular characterization of mucosal adherent bacteria and associations with colorectal adenomas. Gut Microbes 2010, 1:138–147.

12. Turnbaugh PJ, Ridaura VK, Faith JJ, Rey FE, Knight R, Gordon JI: The effect of diet on the human gut microbiome: a metagenomic analysis in humanized gnotobiotic mice. Sci Transl Med 2009, 1:6ra14.

13. Ridaura VK, Faith JJ, Rey FE, Cheng J, Duncan AE, Kau AL, Griffin NW, Lombard V, Henrissat B, Bain JR, Muehlbauer MJ, Ilkayeva O, Semenkovich CF, Funai K, Hayashi DK, Lyle BJ, Martini MC, Ursell LK, Clemente JC, Van Treuren W, Walters WA, Knight R, Newgard CB, Heath AC, Gordon JI: Gut microbiota from twins discordant for obesity modulate metabolism in mice. Science 2013, 341:1241214.

14. Kesse E, Clavel-Chapelon F, Boutron-Ruault MC: Dietary patterns and risk of colorectal tumors: a cohort of French women of the National Education System (E3N). Am J Epidemiol 2006, 164:1085–1093.

15. Huxley RR, Ansary-Moghaddam A, Clifton P, Czernichow S, Parr CL, Woodward M: The impact of dietary and lifestyle risk factors on risk of colorectal cancer: a quantitative overview of the epidemiological evidence. Int J Cancer 2009, 125:171–180.

16. Kune S, Kune G, Watson L: Case-control study of dietary etiological factors: The Melbourne colorectal cancer study. Nutrition and cancer 1987.

17. Schulz MD, Atay C, Heringer J, Romrig FK, Schwitalla S, Aydin B, Ziegler PK, Varga J, Reindl W, Pommerenke C, Salinas-Riester G, Böck A, Alpert C, Blaut M, Polson SC, Brandl L, Kirchner T, Greten FR, Polson SW, Arkan MC: High-fat-diet-mediated dysbiosis promotes intestinal carcinogenesis independently of obesity. Nature 2014, 514:508–512.

18. Goodrich JK, Davenport ER, Beaumont M, Jackson MA, Knight R, Ober C, Spector TD, Bell JT, Clark AG, Ley RE: Genetic determinants of the gut microbiome in UK twins. Cell Host Microbe 2016, 19:731–743.

19. Goodrich JK, Davenport ER, Waters JL, Clark AG, Ley RE: Cross-species comparisons of host genetic associations with the microbiome. Science 2016, 352:532–535.

20. Goodrich JK, Waters JL, Poole AC, Sutter JL, Koren O, Blekhman R, Beaumont M, Van Treuren W, Knight R, Bell JT, Spector TD, Clark AG, Ley RE: Human genetics shape the gut microbiome. Cell 2014, 159:789–799.

21. Blekhman R, Goodrich JK, Huang K, Sun Q, Bukowski R, Bell JT, Spector TD, Keinan A, Ley RE, Gevers D, Clark AG: Host genetic variation impacts microbiome composition across human body sites. Genome Biol 2015, 16:191.

22. Knights D, Silverberg MS, Weersma RK, Gevers D, Dijkstra G, Huang H, Tyler AD, van Sommeren S, Imhann F, Stempak JM, Huang H, Vangay P, Al-Ghalith GA, Russell C, Sauk J, Knight J, Daly MJ, Huttenhower C, Xavier RJ: Complex host genetics influence the microbiome in inflammatory bowel disease. Genome Med 2014, 6:107.

23. Davenport ER, Cusanovich DA, Michelini K, Barreiro LB, Ober C, Gilad Y: Genome-Wide Association Studies of the Human Gut Microbiota. PLoS ONE 2015, 10:e0140301.

24. Cancer Genome Atlas Network: Comprehensive molecular characterization of human colon and rectal cancer. Nature 2012, 487:330–337.

25. Burns MB, Montassier E, Abrahante J, Starr TK, Knights D, Blekhman R: Discrete mutations in colorectal cancer correlate with defined microbial communities in the tumor microenvironment. bioRxiv 2016.

26. Dove WF, Clipson L, Gould KA, Luongo C, Marshall DJ, Moser AR, Newton MA, Jacoby RF: Intestinal neoplasia in the ApcMin mouse: independence from the microbial and natural killer (beige locus) status. Cancer Res 1997, 57:812–814.

27. Uronis JM, Mühlbauer M, Herfarth HH, Rubinas TC, Jones GS, Jobin C: Modulation of the intestinal microbiota alters colitis-associated colorectal cancer susceptibility. PLoS ONE 2009, 4:e6026.

28. Liu S, da Cunha AP, Rezende RM, Cialic R, Wei Z, Bry L, Comstock LE, Gandhi R, Weiner HL: The host shapes the gut microbiota via fecal microrna. Cell Host Microbe 2016, 19:32– 43.

29. Li L, Sarver AL, Khatri R, Hajeri PB, Kamenev I, French AJ, Thibodeau SN, Steer CJ, Subramanian S: Sequential expression of miR-182 and miR-503 cooperatively targets FBXW7, contributing to the malignant transformation of colon adenoma to adenocarcinoma. J Pathol 2014, 234:488–501.

30. Li Y, Lauriola M, Kim D, Francesconi M, D’Uva G, Shibata D, Malafa MP, Yeatman TJ, Coppola D, Solmi R, Cheng JQ: Adenomatous polyposis coli (APC) regulates miR17-92 cluster through β-catenin pathway in colorectal cancer. Oncogene 2016, 35:4558–4568.

31. Diosdado B, van de Wiel MA, Terhaar Sive Droste JS, Mongera S, Postma C, Meijerink WJHJ, Carvalho B, Meijer GA: MiR-17-92 cluster is associated with 13q gain and c-myc expression during colorectal adenoma to adenocarcinoma progression. Br J Cancer 2009, 101:707–714.

32. Hu S, Liu L, Chang EB, Wang J-Y, Raufman J-P: Butyrate inhibits pro-proliferative miR-92a by diminishing c-Myc-induced miR-17-92a cluster transcription in human colon cancer cells. Mol Cancer 2015, 14:180.

33. Peck BCE, Mah AT, Pitman WA, Ding S, Lund PK, Sethupathy P: Functional transcriptomics in diverse intestinal epithelial cell types reveals robust microrna sensitivity in intestinal stem cells to microbial status. J Biol Chem 2017, 292:2586–2600.

34. Cai L, Ye L, Tong AHY, Lok S, Zhang T: Biased diversity metrics revealed by bacterial 16S pyrotags derived from different primer sets. PLoS ONE 2013, 8:e53649.

35. Edgar RC: Search and clustering orders of magnitude faster than BLAST. Bioinformatics 2010, 26:2460–2461.

36. DeSantis TZ, Hugenholtz P, Larsen N, Rojas M, Brodie EL, Keller K, Huber T, Dalevi D, Hu P, Andersen GL: Greengenes, a chimera-checked 16S rRNA gene database and workbench compatible with ARB. Appl Environ Microbiol 2006, 72:5069–5072.

37. Navas-Molina JA, Peralta-Sánchez JM, González A, McMurdie PJ, Vázquez-Baeza Y, Xu Z, Ursell LK, Lauber C, Zhou H, Song SJ, Huntley J, Ackermann GL, Berg-Lyons D, Holmes S, Caporaso JG, Knight R: Advancing our understanding of the human microbiome using QIIME. Meth Enzymol 2013, 531:371–444.

38. Caporaso JG, Kuczynski J, Stombaugh J, Bittinger K, Bushman FD, Costello EK, Fierer N, Peña AG, Goodrich JK, Gordon JI, Huttley GA, Kelley ST, Knights D, Koenig JE, Ley RE, Lozupone CA, McDonald D, Muegge BD, Pirrung M, Reeder J, Sevinsky JR, Turnbaugh PJ, Walters WA, Widmann J, Yatsunenko T, Zaneveld J, Knight R: QIIME allows analysis of high-throughput community sequencing data. Nat Methods 2010, 7:335–336.

39. Bolger AM, Lohse M, Usadel B: Trimmomatic: a flexible trimmer for Illumina sequence data. Bioinformatics 2014, 30:2114–2120.

40. Langmead B, Trapnell C, Pop M, Salzberg SL: Ultrafast and memory-efficient alignment of short DNA sequences to the human genome. Genome Biol 2009, 10:R25.

41. Masella AP, Bartram AK, Truszkowski JM, Brown DG, Neufeld JD: PANDAseq: paired-end assembler for illumina sequences. BMC Bioinformatics 2012, 13:31.

42. Anders S, Pyl PT, Huber W: HTSeq--a Python framework to work with high-throughput sequencing data. Bioinformatics 2015, 31:166–169.

43. Lopez JP, Diallo A, Cruceanu C, Fiori LM, Laboissiere S, Guillet I, Fontaine J, Ragoussis J, Benes V, Turecki G, Ernst C: Biomarker discovery: quantification of microRNAs and other small non-coding RNAs using next generation sequencing. BMC Med Genomics 2015, 8:35.

44. Love MI, Huber W, Anders S: Moderated estimation of fold change and dispersion for RNA-seq data with DESeq2. Genome Biol 2014, 15:550–550.

45. Friedman J, Alm EJ: Inferring correlation networks from genomic survey data. PLoS Comput Biol 2012, 8:e1002687.

46. Langille MGI, Zaneveld J, Caporaso JG, McDonald D, Knights D, Reyes JA, Clemente JC, Burkepile DE, Vega Thurber RL, Knight R, Beiko RG, Huttenhower C: Predictive functional profiling of microbial communities using 16S rRNA marker gene sequences. Nat Biotechnol 2013, 31:814–821.

47. Vlachos IS, Zagganas K, Paraskevopoulou MD, Georgakilas G, Karagkouni D, Vergoulis T, Dalamagas T, Hatzigeorgiou AG: DIANA-miRPath v3.0: deciphering microRNA function with experimental support. Nucleic Acids Res 2015, 43:W460–6.

48. Lewis BP, Burge CB, Bartel DP: Conserved seed pairing, often flanked by adenosines, indicates that thousands of human genes are microRNA targets. Cell 2005, 120:15–20.

49. Sarver AL, French AJ, Borralho PM, Thayanithy V, Oberg AL, Silverstein KAT, Morlan BW, Riska SM, Boardman LA, Cunningham JM, Subramanian S, Wang L, Smyrk TC, Rodrigues CMP, Thibodeau SN, Steer CJ: Human colon cancer profiles show differential microRNA expression depending on mismatch repair status and are characteristic of undifferentiated proliferative states. BMC Cancer 2009, 9:401.

50. Mogilyansky E, Rigoutsos I: The miR-17/92 cluster: a comprehensive update on its genomics, genetics, functions and increasingly important and numerous roles in health and disease. Cell Death Differ 2013, 20:1603–1614.

51. Oberg AL, French AJ, Sarver AL, Subramanian S, Morlan BW, Riska SM, Borralho PM, Cunningham JM, Boardman LA, Wang L, Smyrk TC, Asmann Y, Steer CJ, Thibodeau SN: miRNA expression in colon polyps provides evidence for a multihit model of colon cancer. PLoS ONE 2011, 6:e20465.

52. Weir TL, Manter DK, Sheflin AM, Barnett BA, Heuberger AL, Ryan EP: Stool microbiome and metabolome differences between colorectal cancer patients and healthy adults. PLoS ONE 2013, 8:e70803.

53. Geng J, Fan H, Tang X, Zhai H, Zhang Z: Diversified pattern of the human colorectal cancer microbiome. Gut Pathog 2013, 5:2.

54. Vogtmann E, Hua X, Zeller G, Sunagawa S, Voigt AY, Hercog R, Goedert JJ, Shi J, Bork P, Sinha R: Colorectal Cancer and the Human Gut Microbiome: Reproducibility with Whole-Genome Shotgun Sequencing. PLoS ONE 2016, 11:e0155362.

55. Yu J, Feng Q, Wong SH, Zhang D, Liang QY, Qin Y, Tang L, Zhao H, Stenvang J, Li Y, Wang X, Xu X, Chen N, Wu WKK, Al-Aama J, Nielsen HJ, Kiilerich P, Jensen BAH, Yau TO, Lan Z, Jia H, Li J, Xiao L, Lam TYT, Ng SC, Cheng AS-L, Wong VW-S, Chan FKL, Xu X, Yang H, et al.: Metagenomic analysis of faecal microbiome as a tool towards targeted non-invasive biomarkers for colorectal cancer. Gut 2017, 66:70–78.

56. Dharmani P, Strauss J, Ambrose C, Allen-Vercoe E, Chadee K: Fusobacterium nucleatum infection of colonic cells stimulates MUC2 mucin and tumor necrosis factor alpha. Infect Immun 2011, 79:2597–2607.

57. Abed J, Emgård JEM, Zamir G, Faroja M, Almogy G, Grenov A, Sol A, Naor R, Pikarsky E, Atlan KA, Mellul A, Chaushu S, Manson AL, Earl AM, Ou N, Brennan CA, Garrett WS, Bachrach G: Fap2 Mediates Fusobacterium nucleatum Colorectal Adenocarcinoma Enrichment by Binding to Tumor-Expressed Gal-GalNAc. Cell Host Microbe 2016, 20:215–225.

58. Andrian E, Grenier D, Rouabhia M: Porphyromonas gingivalis gingipains mediate the shedding of syndecan-1 from the surface of gingival epithelial cells. Oral Microbiol Immunol 2006, 21:123–128.

59. Benvenuti S, Sartore-Bianchi A, Di Nicolantonio F, Zanon C, Moroni M, Veronese S, Siena S, Bardelli A: Oncogenic activation of the RAS/RAF signaling pathway impairs the response of metastatic colorectal cancers to anti-epidermal growth factor receptor antibody therapies. Cancer Res 2007, 67:2643–2648.

60. Hynes NE, Lane HA: ERBB receptors and cancer: the complexity of targeted inhibitors. Nat Rev Cancer 2005, 5:341–354.

61. Konsavage WM, Kyler SL, Rennoll SA, Jin G, Yochum GS: Wnt/β-catenin signaling regulates Yes-associated protein (YAP) gene expression in colorectal carcinoma cells. J Biol Chem 2012, 287:11730–11739.

62. De Luca A, Maiello MR, D’Alessio A, Pergameno M, Normanno N: The RAS/RAF/MEK/ERK and the PI3K/AKT signalling pathways: role in cancer pathogenesis and implications for therapeutic approaches. Expert Opin Ther Targets 2012, 16 Suppl 2:S17–27.

63. Tjalsma H, Boleij A, Marchesi JR, Dutilh BE: A bacterial driver-passenger model for colorectal cancer: beyond the usual suspects. Nat Rev Microbiol 2012, 10:575–582.

64. Rubinstein MR, Wang X, Liu W, Hao Y, Cai G, Han YW: Fusobacterium nucleatum promotes colorectal carcinogenesis by modulating E-cadherin/β-catenin signaling via its FadA adhesin. Cell Host Microbe 2013, 14:195–206.

65. Tahara T, Yamamoto E, Suzuki H, Maruyama R, Chung W, Garriga J, Jelinek J, Yamano H, Sugai T, An B, Shureiqi I, Toyota M, Kondo Y, Estécio MRH, Issa J-PJ: Fusobacterium in colonic flora and molecular features of colorectal carcinoma. Cancer Res 2014, 74:1311– 1318.

66. Castellarin M, Warren RL, Freeman JD, Dreolini L, Krzywinski M, Strauss J, Barnes R, Watson P, Allen-Vercoe E, Moore RA, Holt RA: Fusobacterium nucleatum infection is prevalent in human colorectal carcinoma. Genome Res 2012, 22:299–306.

67. Kholodkova E, Kriukov I, Baturo A, et al.: Etiologic role of bacteria of the genus Providencia in acute intestinal diseases. 1977.

68. Shima A, Hinenoya A, Asakura M, Nagita A, et al.: Prevalence of Providencia strains among children with diarrhea in Japan. Japanese journal of 2012.

69. Murata T, Iida T, Shiomi Y, Tagomori K, et al.: A large outbreak of foodborne infection attributed to Providencia alcalifaciens. Journal of Infectious 2001.

70. Wu S, Rhee K-J, Zhang M, Franco A, Sears CL: Bacteroides fragilis toxin stimulates intestinal epithelial cell shedding and gamma-secretase-dependent E-cadherin cleavage. J Cell Sci 2007, 120(Pt 11):1944–1952.

71. Nougayrède J-P, Homburg S, Taieb F, Boury M, Brzuszkiewicz E, Gottschalk G, Buchrieser C, Hacker J, Dobrindt U, Oswald E: Escherichia coli induces DNA double-strand breaks in eukaryotic cells. Science 2006, 313:848–851.

72. Toller IM, Neelsen KJ, Steger M, Hartung ML, Hottiger MO, Stucki M, Kalali B, Gerhard M, Sartori AA, Lopes M, Müller A: Carcinogenic bacterial pathogen Helicobacter pylori triggers DNA double-strand breaks and a DNA damage response in its host cells. Proc Natl Acad Sci U S A 2011, 108:14944–14949.

73. He TC, Sparks AB, Rago C, Hermeking H, Zawel L, da Costa LT, Morin PJ, Vogelstein B, Kinzler KW: Identification of c-MYC as a target of the APC pathway. Science 1998, 281:1509–1512.

74. Mann B, Gelos M, Siedow A, Hanski ML, Gratchev A, Ilyas M, Bodmer WF, Moyer MP, Riecken EO, Buhr HJ, Hanski C: Target genes of beta-catenin-T cell-factor/lymphoid-enhancer-factor signaling in human colorectal carcinomas. Proc Natl Acad Sci U S A 1999, 96:1603–1608.

75. Zhang X, Gaspard JP, Chung DC: Regulation of vascular endothelial growth factor by the Wnt and K-ras pathways in colonic neoplasia. Cancer Res 2001, 61:6050–6054.

76. O’Donnell KA, Wentzel EA, Zeller KI, Dang CV, Mendell JT: c-Myc-regulated microRNAs modulate E2F1 expression. Nature 2005, 435:839–843.

77. Dang CV: c-Myc target genes involved in cell growth, apoptosis, and metabolism. Mol Cell Biol 1999, 19:1–11.

78. Louis P, Hold GL, Flint HJ: The gut microbiota, bacterial metabolites and colorectal cancer. Nat Rev Microbiol 2014, 12:661–672.

79. Baxter NT, Zackular JP, Chen GY, Schloss PD: Structure of the gut microbiome following colonization with human feces determines colonic tumor burden. Microbiome 2014, 2:20.

80. Allali I, Delgado S, Marron PI, Astudillo A, Yeh JJ, Ghazal H, Amzazi S, Keku T, Azcarate-Peril MA: Gut microbiome compositional and functional differences between tumor and non-tumor adjacent tissues from cohorts from the US and Spain. Gut Microbes 2015, 6:161–172.

81. Linden R, Martins VR, Prado MAM, Cammarota M, Izquierdo I, Brentani RR: Physiology of the prion protein. Physiol Rev 2008, 88:673–728.

82. Fang JY, Richardson BC: The MAPK signalling pathways and colorectal cancer. Lancet Oncol 2005, 6:322–327.

83. Yamaguchi H, Bhalla K, Wang H-G: Bax plays a pivotal role in thapsigargin-induced apoptosis of human colon cancer HCT116 cells by controlling Smac/Diablo and Omi/HtrA2 release from mitochondria. Cancer Res 2003, 63:1483–1489.

84. Wynendaele E, Verbeke F, D’Hondt M, Hendrix A, Van De Wiele C, Burvenich C, Peremans K, De Wever O, Bracke M, De Spiegeleer B: Crosstalk between the microbiome and cancer cells by quorum sensing peptides. Peptides 2015, 64:40–48.

85. Tailford LE, Crost EH, Kavanaugh D, Juge N: Mucin glycan foraging in the human gut microbiome. Front Genet 2015, 6:81.

